# ^★^Track: Inferred counting and tracking of replicating DNA loci

**DOI:** 10.1101/2022.12.05.519146

**Authors:** Robin Köhler, Ismath Sadhir, Seán M. Murray

## Abstract

Fluorescent microscopy is the primary method to study DNA organization within cells. However the variability and low signal-to-noise commonly associated with live-cell time lapse imaging challenges quantitative measurements. In particular, obtaining quantitative or mechanistic insight often depends on the accurate tracking of fluorescent particles. Here, we present ^★^Track, an inference method that determines the most likely temporal tracking of replicating intracellular particles such DNA loci while accounting for missing, merged and spurious detections. It allows the accurate prediction of particle copy numbers as well as the timing of replication events. We demonstrate ^★^Track’s abilities and gain new insight into plasmid copy number control and the volume dependence of bacterial chromosome replication initiation. By enabling the accurate tracking of DNA loci, ^★^Track can help to uncover the mechanistic principles of chromosome organisation and dynamics across a range of systems.

**Significance:** Microscopy is one of the main tools for studying the intracellular organisation of cells. In particular, fluorescent proteins allow us to study the dynamics of many cellular processes. However, this requires the accurate tracking of fluorescent foci. Here, we present ^★^Track a tool tailored to the tracking of replicating persistent subcellular particles such as DNA loci. ^★^Track provides accurate predictions of particle copy number and replication timing even in the presence of substantial noise. The knowledge of these quantities are critical for uncovering the mechanisms behind many cell-cycle dependent processes, such the control of chromosome and plasmid replication initiation.

## Introduction

Fluorescence live-cell microscopy is a powerful tool for the study of subcellular organisation and dynamics. However, quantitative analysis requires accurate tracking of the detected fluorescent foci, which may visualize organelles, molecular complexes or even single molecules. This is a challenging task due to high foci density, heterogeneous motion, foci appearance and disappearance and apparent or actual merging and splitting events. Since obtaining the global optimal tracking is computational prohibitive, several heuristic algorithms have been developed to find approximate solutions with greater computational efficiency^1–3^, some of which have been released as publicly available software^4–9^. However, no single algorithm excels at all applications^10^.

Under normal circumstances, chromosomal loci or extrachromosomal DNA, such as plasmids, only increase in number during the cell cycle, typically by a factor of two. This additional information places a significant constraint that can be used to optimize their tracking. At the same time, the typically low copy numbers involved mean that the problem lies at the other extreme of the efficiency-accuracy tradeoff i.e. some of the computationally necessary simplifications required at higher numbers can be avoided. To our knowledge no tracking software is tailored to this scenario and we found that existing tools did not perform with sufficiently high accuracy. Note that despite the low copy numbers, generating the optimal tracking is not trivial since false positive (i.e. spurious) and false negative detections (due to the foci moving out of the focal plane, temporary merging or detection failure) can make it challenging to determine the time point at which foci are duplicated even at relatively low copy numbers. This is especially true for bacteria due their small size and the smaller pool of fluorescent protein.

Here, we present ^★^Track (pronounced ‘star-track’)^1^ a tool for the accurate tracking of replicating chromosomal and extrachromosomal DNA, and other low-copy number persistent particles, from time lapse images of live cells. The algorithm is transparent and user friendly and is completely specified, in the default case, by only four input parameters (there are no hidden internal parameters). Matlab and Python implementations are provided so that ^★^Track can be easily integrated into existing spot detection pipelines.

## Materials and Methods

### Strains and growth condition

F plasmid experiments use strain DLT3125^11^, a derivative of the E. coli K-12 strain DLT1215^12^ containing the mini-F plasmid derivative pJYB234. This plasmid carries a functional ParB-mVenus fusion. Overnight cultures were grown at 37°C in LB-Media supplemented with 10 μg/ml thymine + 10 μg/ml chloramphenicol. The strain IS130 was constructed by transduction of matP-YPet from RH3^13^ and oriC-*parS*_P1_ from strain RM3^14,15^ into Escherichia coli K-12 MG1655 strain (lab collection) which was then transformed with pFHCP1-mTurquoise2 plasmid. The plasmid pFHCP1-mTurquoise2 was derived from plasmid pFHC2973 by deletion of ygfp-parB_pMT1_ and replacement of CFP with mTurquoise2^14,16^. Overnight cultures were grown in M9 minimal media supplemented with 0.2% glucose, 2 mM MgSO_4_, 0.1 mM CaCl_2_ and 0.5 mg/mL BSA.

### Microfluidics

Like the original mother machine^17^, our design consists of a main channel through which nutrient media flows and narrow growth-channels in which cells are trapped. However, we follow Baltekin et al.^18^ and include i) a small opening at the end of each growth channel ii) a waste channel connected to that opening to allow a continuous flow of nutrients through the growth channels iii) an inverted growth-channel that is used to remove the background from fluorescence and phase contrast. Before imaging, the chip is bound to a glass slide using a plasma generator and baked for 30 minutes at 80°C.

### Microscopy

We used a Nikon Ti microscope with a 100x/1.45 oil objective and a Hamamatsu Photonics camera for all imaging. For imaging cells of strain DLT3125 and IS130 we used a mother machine. Overnight cultures of DLT3125 were inoculated into fresh media (M9 + 0.5% glycerol + 0.2% casamino acids + 0.04 mg/mL thymine + 0.2 mg/mL leucine + 10 μg/mL chloramphenicol) for 4 hours at 30°C before imaging. For the strain IS130, 50 μM IPTG (for induction of mTurquoise2-ParB_P1_) was added to the media defined in the ‘Strains and growth condition’ section 2 hours before and during the experiment. Cells were loaded into the chip through the main channel and the chip was placed into a preheated microscope at 30°C. The cells were constantly supplied with fresh media by pumping 2 μl per minute through the microfluidic chip. Cells were grown for at least 2 hours inside the microscope before imaging. DLT3125 was imaged at 1 minute intervals and IS130 was imaged at 5 minute intervals for approximately 72 hours. Both phase contrast and fluorescent signal were captured.

### Image processing

Our image processing pipeline, Mothersegger (https://gitlab.gwdg.de/murray-group/MotherSegger) has been described previously^19^. Briefly, it consists of four parts: I) preprocessing, II) segmentation III) cell tracking and IV) foci detection. Parts I, III and IV use custom Matlab scripts, while Part II is based on SuperSegger^20^, a Matlab-based package for segmenting and tracking bacteria within microcolonies, that we modified to better handle high-throughput data. In Part I each frame of an acquired image stack is aligned (the offset between frames in x and y is removed). Afterwards the image stack is rotated so the growth channels are vertical. A mask of the mother machine layout is fitted to the phase contrast, using cross-correlation, to identify where the growth channels are located. Each growth channel is extracted from the image stack and the flipped inverted channel is subtracted to remove the background from both the fluorescence signal and phase contrast. In Part III the cells are tracked. Since cells cannot change their order inside the growth channel, they can be tracked by matching similar cell length between frames (starting from the bottom of the channel). Cell cycles that do not have exactly 1 parent and 2 daughters are excluded from analysis along with their immediate relatives (with the exception of who are pushed out of the growth channel). Part IV detects fluorescent foci within cells. It is based on the SpotFinderZ tool from Microbetracker^21^.

### Computer generated trajectories and manipulation

The trajectories used in Fig. 1c-f were generated with our previous published model of plasmid positioning^19^. This is a stochastic model of plasmid positioning by the interaction of plasmid-bound ParB with nucleoid associated ParA. These trajectories were used as ground truth. As stated in Fig. 1 we manipulated the ground truth by adding (false positives) and removing (false negatives) a percentage of foci. If the ground truth contains 100 foci and 10% false positives are added and 5% false negatives are removed the resulting modified data set contains 95 foci (100 - 10 fn + 5 fp).

**Fig. 1:**
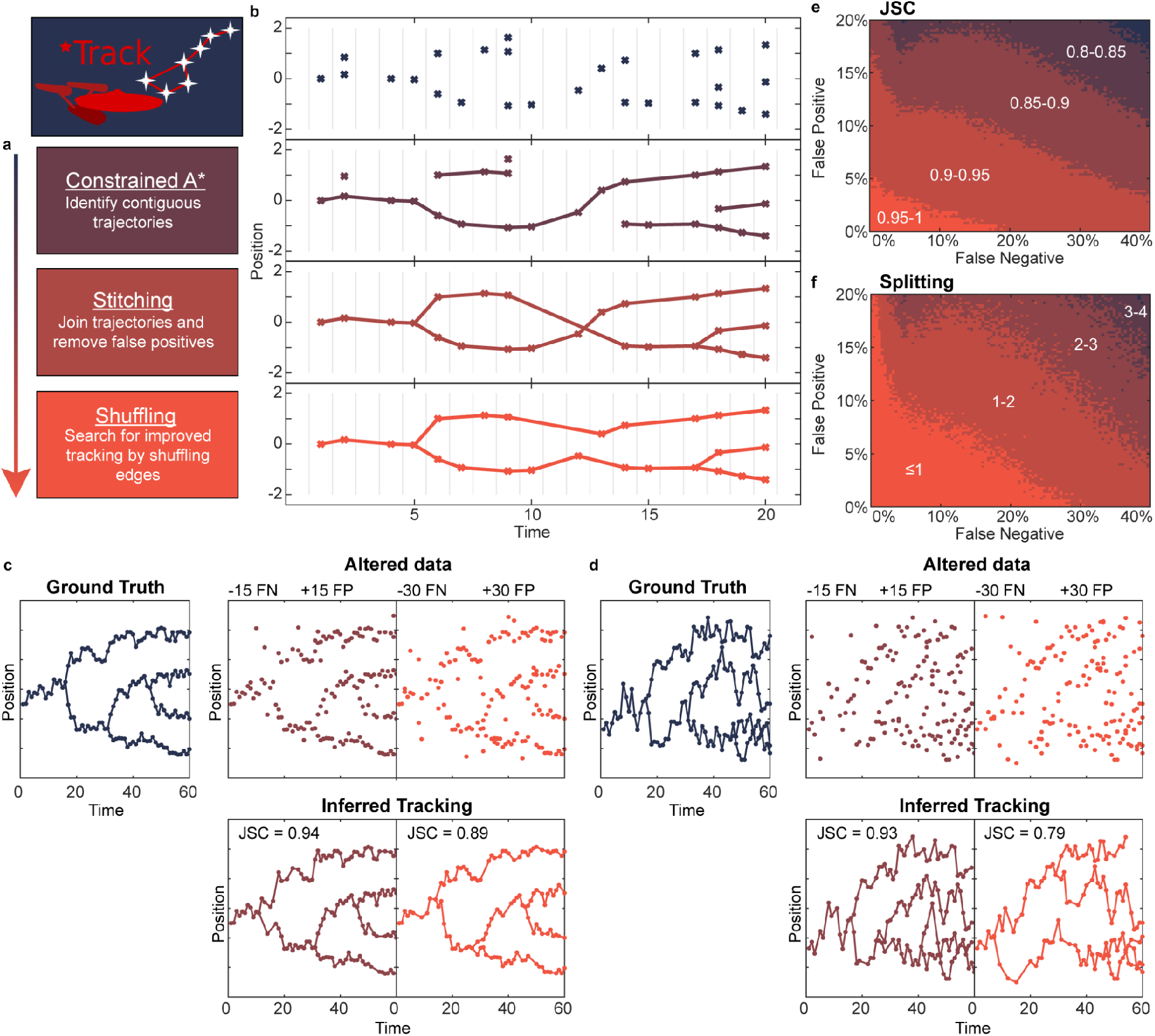
Overview and accuracy of ^★^Track. **a**, All main steps of the algorithm with a short description. The arrow indicates the order. **b**, Example timelapse of localisations and how the data looks after each step in **a. c**, A simulated trajectory (150 foci) is used as ground truth to measure the accuracy of ^★^Track. The ground truth is altered by adding a number of spurious foci (false positive, FP) and removing a number of real foci (false negative, FN) (panels labeled Altered data). ^★^Track was then used to infer a tracking of the altered data (panels labeled Inferred). Accuracy is quantified using the Jaccard similarity coefficient (JSC).The JSC is equal to the number of real foci included in the tracking divided by the sum of the number of spurious foci in the tracking and the total number of real foci in the altered data. The JSC is between 0 and 1, where 1 is a perfect score (the tracking contains all the real and no spurious foci) and 0 is the worst score (no real foci included in the tracking). **d**, Same as **c** but for more mobile particles. **e**, JSC for a wide range of false positive and false negative rates. At each point, 100 trajectories were generated as in **c. f**, Same as in **e** but for splitting accuracy. The number indicates the mean absolute deviation of the splitting event in frames from the ground truth.

### A* algorithm

The first step of ^★^Track is based on the A* graph-traversal algorithm, which we first summarize. A graph consists of nodes connected by edges that have a weight corresponding to the cost of traversing that edge. The cost of a path on the graph is the cumulative cost of all its edges. Given a start and target node of a graph, the A*-algorithm finds the shortest path between them. A path is considered the shortest if there exists no other path connecting these two nodes with a lower cost. The algorithm begins by initializing a candidate list of partial paths consisting of only the first edge (the edges of the starting node). The key feature of the algorithm is the generation of a lower bound for the cost of completing a partial path. The determination of this bound is problem specific. The algorithm then selects the first edge that gives the lowest lower bound for the total cost (the cost of the first edge plus the lower bound for completing the path starting from that edge). In the second iteration, all possible choices of second edge (with the chosen first edge) are added to the candidate list and again the best second edge is selected based on the cost estimate. This process repeats until either the target node is reached or the estimated cost of the current best partial path exceeds the estimated cost of completing a partial path further up the graph, e.g. a path with a different first edge. In this case, the process then continues from this different first edge. In this way the algorithm eventually finds the path with global minimum cost.

### Cost of linking two foci

Use of the A* algorithm requires the specification of a cost for linking focus *i* on one frame to focus *j* on a later frame. For this, we assume that foci move diffusively between frames and consider the probability that the 2D distance traveled would be at least as great as that observed

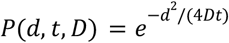

where *d* is the 2D spatial distance between the foci, t is the number of frames between foci and *D* is the diffusion coefficient. The movement cost is then *c*_*i,j*_ = – *log*(*P*(*d, t, D*)). An alternative model can be used if required. If the two foci are not from consecutive frames then we add a cost *c_neg_* = – *loq*(*f_neg_*) for each of the intermediate *t* – 1 ≤ *m* frames, where *f_neg_* is the probability of a false negative and *m* is a user-provided upper bound for the number of frames on which the focus was missing.

### Layered Graph

The second requirement of the A* algorithm is that the search is between start and target nodes. On the other hand, the initial tracking consists of time-directed connections between foci that need not form a connected path. We therefore developed a mapping of the temporal tracking problem to a radial layered graph structure on which the A* algorithm could be applied.

We illustrate this with an example in Fig. S2. Panel S2a shows an example data set consisting of *n_foci_* = 6 foci localisations across *t_end_* = 5 frames. We first determine the set of acceptable links between these foci. Acceptable links are those with a cost below the threshold *c_th_* = *min*(– *m log*(*f_neg_*), *cmax*) based on the user provided maximum number of frame skips, *m* and the false negative probability. This threshold ensures no link is allowed between foci separated by more than *m* intermediate frames, as well as very spatially distant links across fewer frames. We also include an upper bound on the threshold *cmax* = – *log*(0. 0001) to account for when *f_neg_* = 0 (no false negatives allowed) in which case the threshold would otherwise be infinite.

Given these acceptable links, we can then create the layered graph in Fig. S2b. This graph contains one layer for each focus plus an additional outermost layer to determine when the tracking is complete. Every path from the first to the last layer corresponds to one possible tracking of the foci in Fig. S2a. The relationship to a tracking is as follows. An edge from **layer** *i* into a numbered **node** *j* in layer *i* + 1 corresponds to an acceptable link between focus *i* and focus *j*. An outgoing edge into a node labelled ‘x’ corresponds to focus *i* having no outgoing link. Note that the layer into which an edge goes, does not matter for the interpretation. A path from layer 1 to the outermost layer crosses each layer once, therefore each layer (except the outermost) has exactly one outgoing edge. This ensures that each focus has at most 1 outgoing link. Furthermore each path contains each numbered node at most once. If a path traverses a numbered node *i*, node *i* will not appear again in the higher layers. This ensures that each focus has only one incoming link. The construction of the graph is illustrated in Supplementary Video 1.

Each edge in the graph has a weight (referred to but not shown in Fig. S2b) which corresponds to the cost of linking the foci associated with that edge, as described above. An edge going from layer *i* into a numbered node *j* has a cost *c_i,j_*. An edge from layer *i* going into a node labelled *‘x’* has a cost equal to 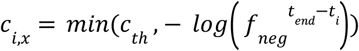 where *t_end_* – *t_i_* is the number of frames between that of focus *i*, *t_i_* and the last frame *t_end_*. The cost of a path is the cumulative cost of its edges and the optimal tracking is the path from the root node in layer 1 to the outermost layer with the lowest cost.

To use the A* algorithm we require a heuristic lower-bound estimate of the cost of reaching the outermost layer from each node in the graph. A path from layer *i* has to traverse all subsequent layers to reach the outermost layer. A lower bound for the cost is then the sum of all the lowest costing outgoing links 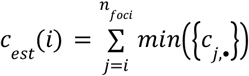 where the minimum is over the cost of all acceptable outgoing links from layer (focus) *j* (including to the ‘*x*’ node).

Since the number of nodes in each layer increases exponentially, it is infeasible, and also unnecessary, to pre-generate the entire graph when tracking more than a few foci over a small number of frames. Therefore the graph is generated dynamically as the A* algorithm traverses it. An example can be seen in Supplementary Video 2. Note that the optimal tracking is found without generating the entire graph. To further reduce the complexity, the candidate list is filtered once a new layer is reached. All candidate paths ending 1 + *max*({*j* – *i* : *t_j_* – *t_i_* ≤ *m*}) layers below the newly reached layer are removed from the candidate list. Therefore, the greater the number *m* of allowed frame skips (consecutive false negatives), the more of the graph is retained. Additionally, if the candidate list exceeds 10^6^ entries, the 10% of partial paths with the highest cost estimates are removed. This is why we refer to the algorithm as constrained.

### Stitching

The goal of this procedure is to integrate the trajectory segments returned by the A* algorithm such that the number of foci monotonically increases. This is done in 3 steps. First, loose beginnings (foci which have no incoming link) are linked to earlier foci (Fig. S3b), either to a real focus according to the lowest cost of the link or to an imaginary focus before the first frame. The latter is for implementation reasons only.

In the next step, loose ends (foci which have no outgoing link) are integrated into the tracking (Fig. S3d). This occurs in three different ways: (1) a loose end can be linked to an imaginary focus after the last frame (Fig. S3c ii & iii), impling some number of missed foci (false negatives) when the loose end is not on the last frame; (2) all the foci of the segment containing the loose end up to any branching point are considered as false positives; (3) a loose end can be interwoven into the tracking by replacing an existing link, which can but does not have to be a link created by integrating a loose beginning (Fig. S3c iv & v). This may create a different loose end, which is treated as in (2). Which method occurs is determined by the cost of the entire tracking.

The cost of the tracking is calculated differently than in the A* step as it incorporates the concept of false positives and applies to the entire tracking and not just disconnected segments. Furthermore, the threshold cost of the A* step is no longer used. Each false positive focus has a cost *c_pos_* = –*log*(*f_pos_*) where *f_pos_* is the false positive probability. Each real focus then has a cost arising from its probability to not be a false positive *c_real_* = – *log*(1 – *f_pos_*). The cost of a tracking is then 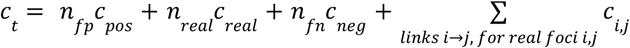, where *n_fp_* is the number of false positives, *n_real_* is the number of detected foci and *n_fn_* is the number of false negative foci.

Returning to the integration of loose ends, the change in cost of the tracking is calculated for every possible way to integrate a loose end and the method with the lowest cost is chosen. The procedure is iterative starting at the last loose end. It continues until there are no loose ends left or the remaining loose ends consist of false positives. In the last step the foci identified as false positives are removed from the tracking (Fig. S3e).

### Shuffling

Shuffling resolves remaining inaccuracies in the tracking after the stitching procedure. Inaccuracies can appear due to the constrained nature of the A*-algorithm and the single step nature of the stitching step. Shuffling two links can resolve inaccuracies. Two links connecting four foci, focus *i* linked with focus *j* and focus *n* linked with focus *m*, have a combined movement cost of *c_i,j_* + *c_n,m_*. The links between the foci can be shuffled, subject to *max*(*i,m*) < *min*(*j,n*), such that focus *i* is linked with focus *m* and focus *n* is linked with focus *j*. If the new cost of *c_i,m_* + *c_n,j_* is lower than *c_i,j_* + *c_n,m_* then the new configuration is accepted. Foci are shuffled until there are no pairs of links left that can be shuffled to reduce the cost of the tracking. See Fig. 1b bottom.

## Results and Discussion

^★^Track operates on a provided list of foci localisations (*t, x, y*), consisting of the frame number and location of the fluorescent foci within a cell or region of interest (corrected for movement and growth as required). The algorithm requires a cost function that specifies, for any given focus, how likely it is that another focus on the next or some later frame is the next detection of the same underlying particle (a higher likelihood results in a lower cost). The latter case implies that the same focus was not detected on some intermediate frames and ^★^Track requires the user to specify the maximum number of consecutive frames for which this is allowed to occur (‘not detected’ includes foci that have merged to within the diffraction limit). Having such a cut-off greatly increases the computational efficiency of the first step of the algorithm. This condition is relaxed for subsequent steps.

The default cost function is relatively simple and uses only the positions of foci. However, it could easily be modified to incorporate the spot intensity, goodness of fit or specific models of movement. We have found for the applications studied that spot intensity and shape can vary considerably between frames due to stochastic variation and movement out of the focal plane and we therefore base the cost function only on the foci positions. The default function assumes that foci move diffusively between frames (this is always true on a short enough timescale) and requires only the diffusion coefficient *D*. The user must also specify *p_fn_*, the probability of a spot not being detected on a given frame (false negative) and *p_fp_*, the probability of a detected spot being spurious (false positive). These three parameters can be estimated by analyzing cells with a sufficiently low number of foci and no splitting events (Fig. S1). Precise values are not required as the algorithm is robust with respect to these parameters.

The pipeline consists of three steps (Fig. 1a). First, the tracking problem is formulated as a pathfinding problem on a layered graph (Fig. S2). A constrained A* algorithm then uses the cost function and the max number of frame skips to find the optimal path on this graph and thereby generate candidate trajectory segments (Fig. 1b second panel, Fig. S2). The resulting trajectories are then stitched or interwoven together in such a way that the total cost of the tracking is minimised. Afterwards, each focus is either marked as a false positive and removed or is part of a trajectory that can be traced from the first frame to the last (Fig. 1b third panel, Fig. S3). Finally, a lower cost tracking is searched for by shuffling links in the tracking (Fig. 1b bottom). This helps to mitigate the constrained nature of the first step. Note that ^★^Track does not predict the location of missing foci, but simply their existence. A detailed description of the constrained A* algorithm and the stitching and shuffling steps can be found in the Methods section.

To assess the accuracy of ^★^Track, we generated ground truth data using a stochastic model of plasmid positioning^19^ and mimicked the presence of false positive and false negative foci by randomly adding and removing data points respectively. We found that even after removing a substantial fraction of localisations and adding many false positives, ^★^Track could accurately reproduce the ground truth tracking including correctly identifying the timepoints of plasmid replication (Fig. 1 c,d). Performing a sweep over a range of false positive and false negative rates, we found that ^★^Track performs with consistently high precision (Fig. 1e,f) and outperformed the state of the art and widely employed method u-track^4,5,22^ (Fig. S4).

To test ^★^Track on real data, we applied it to the study of plasmid copy number control. This has been a topic of active research for several decades^23–27^ but progress has more recently slowed due, in part, to the inability to accurately measure plasmid copy numbers in individual cells^28^. Having recently performed a high-throughput study of the partitioning mechanism of the low copy number F plasmid using a microfluidic ‘mother machine’ device^19^, we decided to revisit our data in the context of copy number control. F plasmid is a tractable system in this regard since duplicated plasmids segregate beyond the diffraction limit within about a minute of replication^29^,^30^. We expected that ^★^Track should therefore be able to accurately determine the temporal changes in copy number during the cell cycle.

In Fig. 2a we present an example timelapse showing how the number of detected foci (of plasmid-labeling ParB-mVenus) changes substantially from frame to frame (see Fig. S5 for more examples). As discussed above this can occur due to foci moving out of the focal plane, merging together or simply due to stochastic fluctuations in the number of plasmid-bound ParB-mVenus. However, ^★^Track inferred a consistent tracking that produces a stepwise increasing copy number and predicts the number and frame of the replication events (Fig. 2a). We then applied the algorithm to the entire data set of 4096 cell cycles and examined how the copy number changes between birth and division. The raw data displayed several irregularities such as cells having no detected plasmids at birth and cell cycles in which the number of plasmids decreased (Fig. 2b). All of these inconsistencies were removed by ^★^Track (Fig. 2c), which also inferred a mean copy number at birth ~12% greater than that of the mean number of detected foci. We found a decreasing linear relationship between the number of plasmids gained during the cell cycle and the number at birth (Fig. 2d), indicating a ‘sizer’-like mechanism for copy control^31^. Interestingly, the mean number of plasmids gained did not depend on the growth rate of the individual cell (Fig. 2e), indicating that plasmid production is coupled to the growth rate of the host so as to produce the same number of plasmids irrespective of the cycle duration.

**Fig. 2:**
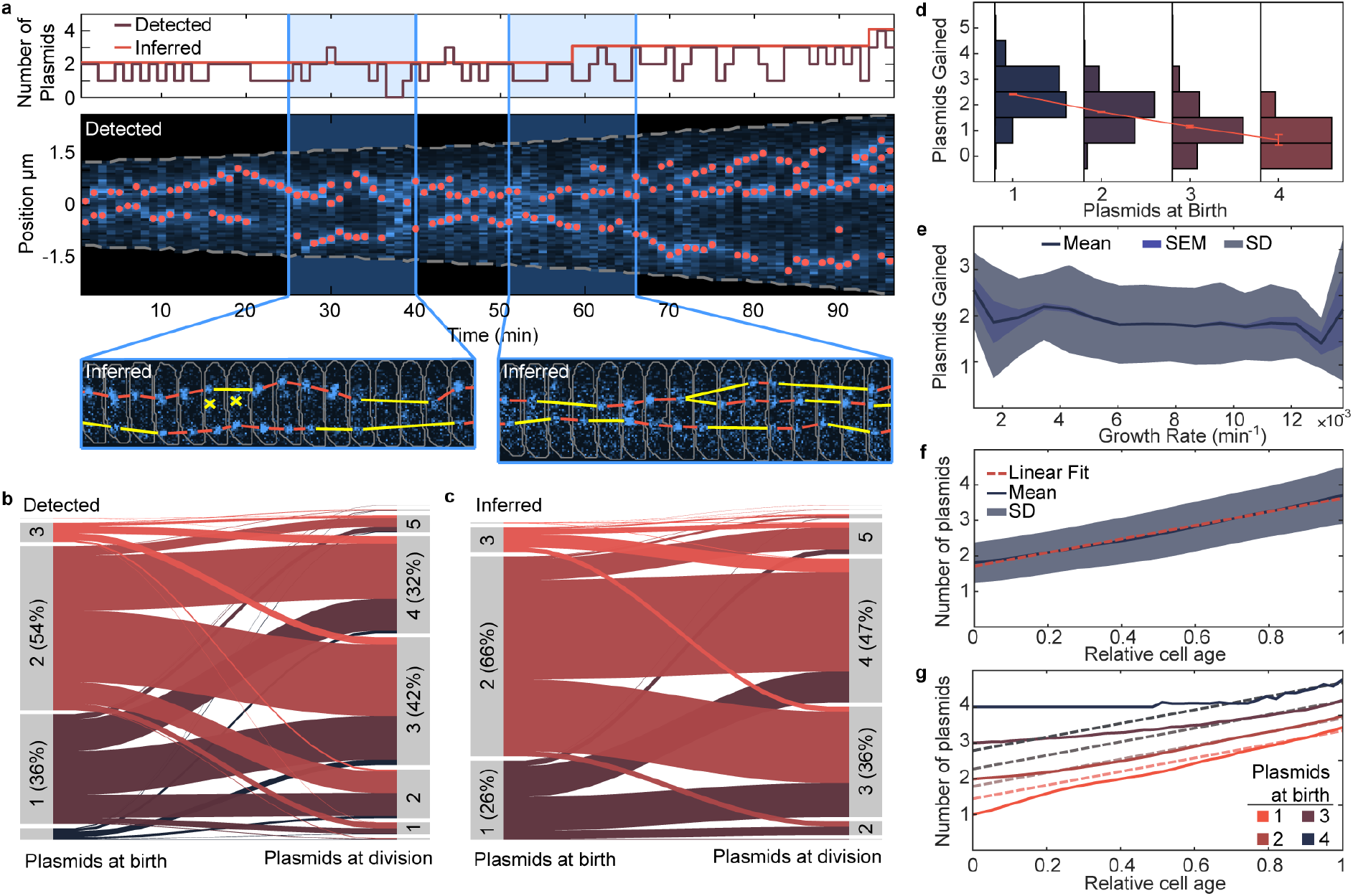
Tracking F-plasmid. **a**, Example cell cycle of the *E. coli* strain DLT3125, hosting an F-plasmid encoding a mVenus-ParB fusion which binds and labels the plasmid. Top: detected and inferred number of plasmids during the cell cycle. Middle: Kymograph of the fluorescent signal of the example cell with detected plasmid locations (red dots). Gray dashed lines indicated cell boundaries. Bottom: Detailed view of the indicated regions. Red and yellow lines indicate the inferred tracking. Yellow lines indicate frames in which missing foci were inferred (false negatives). Red cross indicates foci identified as false positives. Gray lines indicate cell contours. **b**, Alluvial plot showing how the number of plasmids changes from birth to division in the raw data. Frame rate is 1 min^−1^. **c**, Same as **b** but with the inferred data. **d**, Side-ways histograms of plasmids gained during a cell cycle plotted against the number of plasmids at birth. The redline depicts the mean number of plasmids gained for each group ± the standard error of the mean (SEM). **e,** plasmids gained during a cell cycle plotted against the growth rate ± SEM and standard deviation (SD). **f**, Mean number of plasmids ± SD plotted against relative cell age (0 is birth and 1 is division). The red dashed line is a linear fit. **g**, The mean number of plasmids grouped by the number of plasmids at birth plotted against relative cell age. The dashed lines are linear fits with the same slope as in **f** with the intercept chosen by fitting to the last portion of each group.

Going deeper, ^★^Track allows us to probe when within the cell cycle plasmids are replicated. We found that the mean number exhibits a surprising linear relationship with time (Fig. 2f). This is consistent with the net replication rate being constant in time i.e. plasmid are produced at the same rate irrespective of how many are present. However, the explicit nature of the sizer regulation could be seen by binning cell cycles according to the number of plasmids at birth (Fig. 2g). Cells born with fewer/more plasmid than average have a higher/lower net plasmid replication rate initially before returning to the population mean production rate towards the end of the cell cycle. This results in outliers converging rapidly to the mean. While negative feedback has long been known to underlie plasmid copy number control^23–27^, we believe that this is the first time it has been characterized at the level of the cell cycle. Furthermore, our result, that the effect of the regulation is to push the system back to a constant net replication rate, can now be used to test models of copy number control and motivate further study.

Next we used ^★^Track to analyse the replication of the origin region of the *Escherichia coli* chromosome. We used a strain in which the origin of replication (*ori*) is visualized through the P1 *parS*/ParB labeling system using an mTurquoise2-ParB fusion and imaged several thousand cell cycles using the same mother machine device as we used for studying F plasmid. As for that case, foci are not always visible or detected and spurious foci can confound interpretation (Fig. 3a, see Fig. S5b for further examples). This makes it challenging to determine the time of duplicated *ori* separation with certainty. Recent studies have used a replisome reporter to identify initiation events^32,33^, which occur approximately 15 min before separation of *ori* foci^34^; however similar detection issues occur. These irregularities could be seen by comparing the number of *ori* on the first and last frames of the cell cycle (Fig. 3b). Significant cell populations have no detected foci at birth or only one focus at division. However, ^★^Track corrected almost all these inconsistencies (Fig. 3c).

**Fig. 3:**
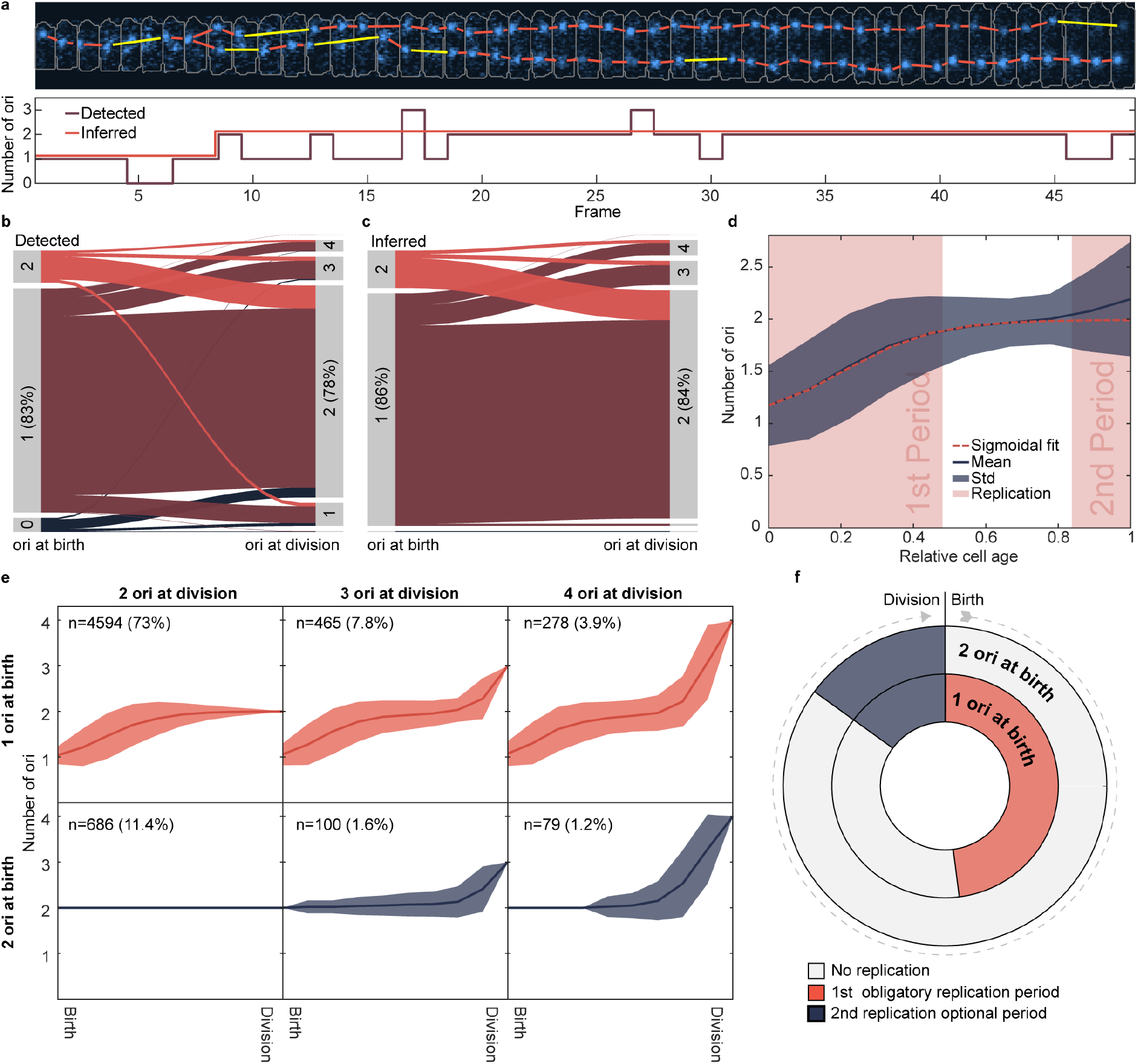
Replication of *ori* occurs during two periods in the cell cycle. **a**, An example cell cycle of the *E. coli* strain IS130. It encodes mTourquise2-ParB which labels the origin of replication (*ori*). Top: Fluorescent signal with tracked *ori*. Bottom: Number of *ori*, both detected and inferred. Frame interval is 5 min. **b**, Alluvial plot showing how the number of *ori* changes from birth to division of the detected foci. **c**, Same as **b** but after ^★^Track was used. **d**, Mean number of *ori* ± standard deviation plotted against relative cell age (0 is birth and 1 is division). The two areas shaded in light red indicate periods during the cell cycle in which *ori* is replicated. The red dashed line is a sigmoidal fit from 0 to 0.85 (excluding the second period). See also Fig. S6. **e**, Data from **d** grouped by number of *ori* at birth (rows) and number of *ori* at division (columns). **f**, Cartoon of replication periods. The plots in **b-e** were created from 6265 cell cycles.

Surprisingly, given the slow-growth conditions used (median doubling time of 140 min), we found that 14% of cells were born with two *ori* (this is a lower bound since a single focus could consist of two unsegregated *ori*). Plotting the number of *ori* against cell age revealed two periods of the cell cycle during which *ori* separation, and hence, replication initiation occurs (Fig. 3d and Fig. 3e). The majority population, cells born with a single *ori* focus, replicated their origin in the first third of the cell cycle. However in about 15% of these cells additional replication events occurred toward the end of the cell cycle producing pre-divisional cells with three or four *ori* foci (Fig. 3e,f). Cells born with two foci only initiated replication in this later period and never at the beginning of the cell cycle, though the majority of such cells do not initiate replication at all and so produce daughter cells with one *ori* focus each.

While analyzing replication initiation in terms of relative cell age is useful, the prevailing understanding is that chromosome replication initiates at an invariant cell volume per *ori*^32,33,35^. Consistent with this, we found that *ori* focus duplication in cells with a single *ori* occured at a cell volume of about 0.8 μm^3^, whereas in cells with two *ori* it occurred at twice this volume (Fig S7). As such, there is really only one replication period defined by the volume per *ori*, but this period overlaps the division event and therefore we are able to observe two different values for the volume at initiation in the same population. This supports previous evidence for the invariant volume per *ori* hypothesis, which has been largely based on comparing populations under different growth conditions.

^★^Track is a powerful new tool for the tracking of plasmid and chromosomal DNA loci. It provides predictions for the timing of replication (initial foci splitting) events that should provide mechanistic insight across a range of systems, and not only for bacteria. ^★^Track can also be applied to any persistent particles that increase in copy number during the cell cycle and could therefore be used to study protein complexes that display this behavior^36–39^. The algorithm is deterministic (there are no stochastic or deep learning components) and therefore produces reproducible results with the user having to specify only four estimatable parameters. ^★^Track is available at https://gitlab.gwdg.de/murray-group/StarTrack.

## Author Contributions

R.K. and S.M.M. designed the research; R.K. developed, tested and applied the method; I.S. performed origin tracking experiments; R.K and S.M.M wrote the paper.

## Acknowledgements

We thank Frédéric Boccard for the strain RM3 and the plasmid pFHC2973, Jaan Mannik for strain RH3 and Jean-Yves Bouet for strain DLT1215 and the plasmid pJYB234.

## Supplementary Figures

**Fig. S1:**
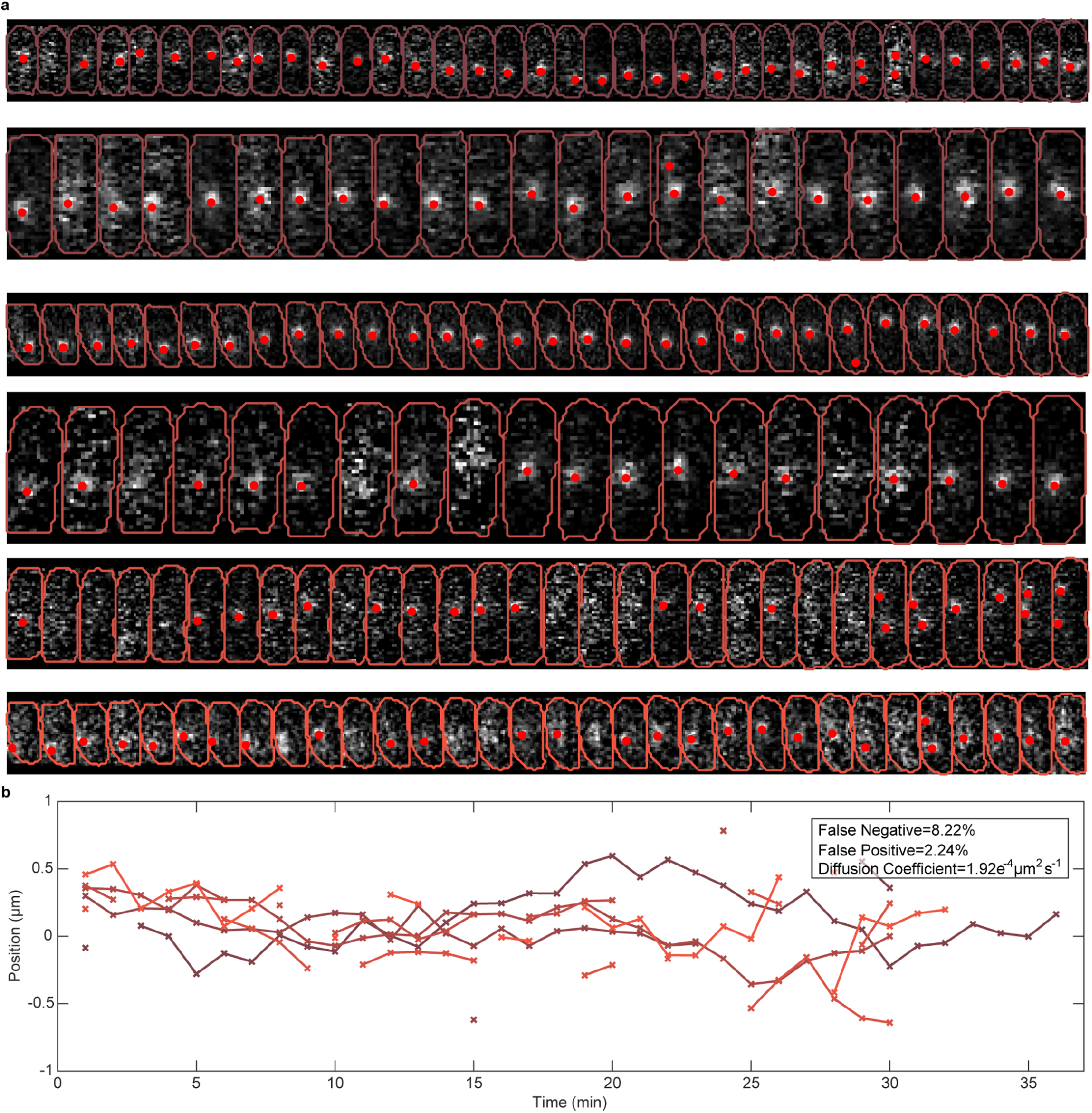
Estimating parameters from data with a low density of foci. **a**, Example cell cycle fragments of the *E. coli* strain DLT3145 containing, except on some isolated frames, a single focus of ParB-mVenus (labeling F plasmid). Cell contours are on top of the fluorescent signal and detected foci are highlighted by red dots. Frame rate is 1 min^−1^. **b**, Foci trajectories from the above cycle fragments. The trajectories contain 9 false positives and 21 false negatives. This is easy to determine by looking at the number of foci on each frame. We found 604 analysable cycle fragments. They contained 361 false positives and 1323 false negatives in a total of 16103 foci. The resulting estimates are a false negative occurrence of 8.22% and a false positive occurrence of 2.24%. Further we estimate the diffusion coefficient of 1.92 x 10^−4^ μm^2^s^−1^ by the variance of the displacement along the long axis between adjacent frames containing only one focus (the step-size distribution) divided by twice the time step.

**Fig. S2:**
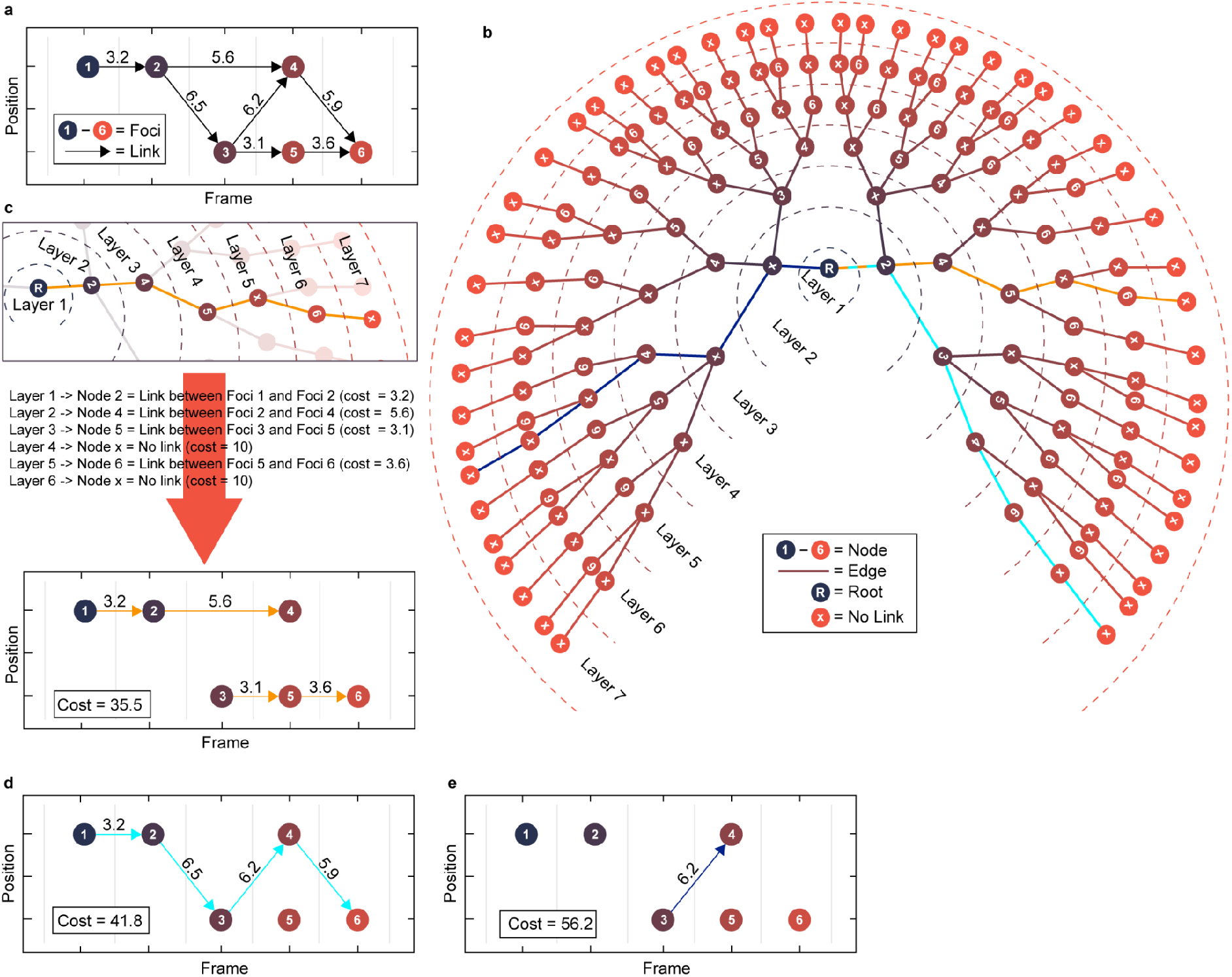
Layered Graph. **a**, Series of 6 foci over 5 frames. ^★^Track first generates a set of acceptable links from every focus to any other focus on a subsequent frame. Acceptable links are those with a cost below a specified threshold, in this case 10 (indicated by arrows), which is obtained from the cost function and the maximum number of skipped frames. **b**, The layered graph generated from **a**. Each direct path from layer 1 to layer 7 corresponds to a unique tracking of the foci in **a**, such that each focus has at most 1 incoming and outgoing link. The graph contains all such trackings. An edge from any node in **layer** *i* to a numbered **node** *j* in the next layer corresponds to a link between focus *i* and focus *j*. An edge from any node in **layer** *i* to a **node** labelled ‘x’ in the next layer corresponds to focus *i* having no outgoing link. Note that it is the layer, not the node, from which the edge originates that determines the meaning of the edge. Each edge has a weight (not shown here) which is equal to the cost of linking the two foci (for edges into numbered nodes) or the cost of a focus not having an outgoing link (for edges into the ‘x’ node). In the latter case, the cost is equal to the threshold mentioned in **a** or if lower the cost of going to the end (see materials). The A*-algorithm generates this graph dynamically as it searches for the optimal tracking. **c**, Example of how to interpret a path in the graph from **b** and how to translate it into trajectory segments. Top: Example path, highlighted in yellow from **b**. Bottom: Resulting trajectory segments with the cumulative cost of all edges of the path. The threshold cost (10) is associated to both foci 4 and 6 for not having an outgoing link. **d&e**, trajectories corresponding to two other paths from **b** as in **c.** The color of the links corresponds to the path in **b** with the same color.

**Fig. S3:**
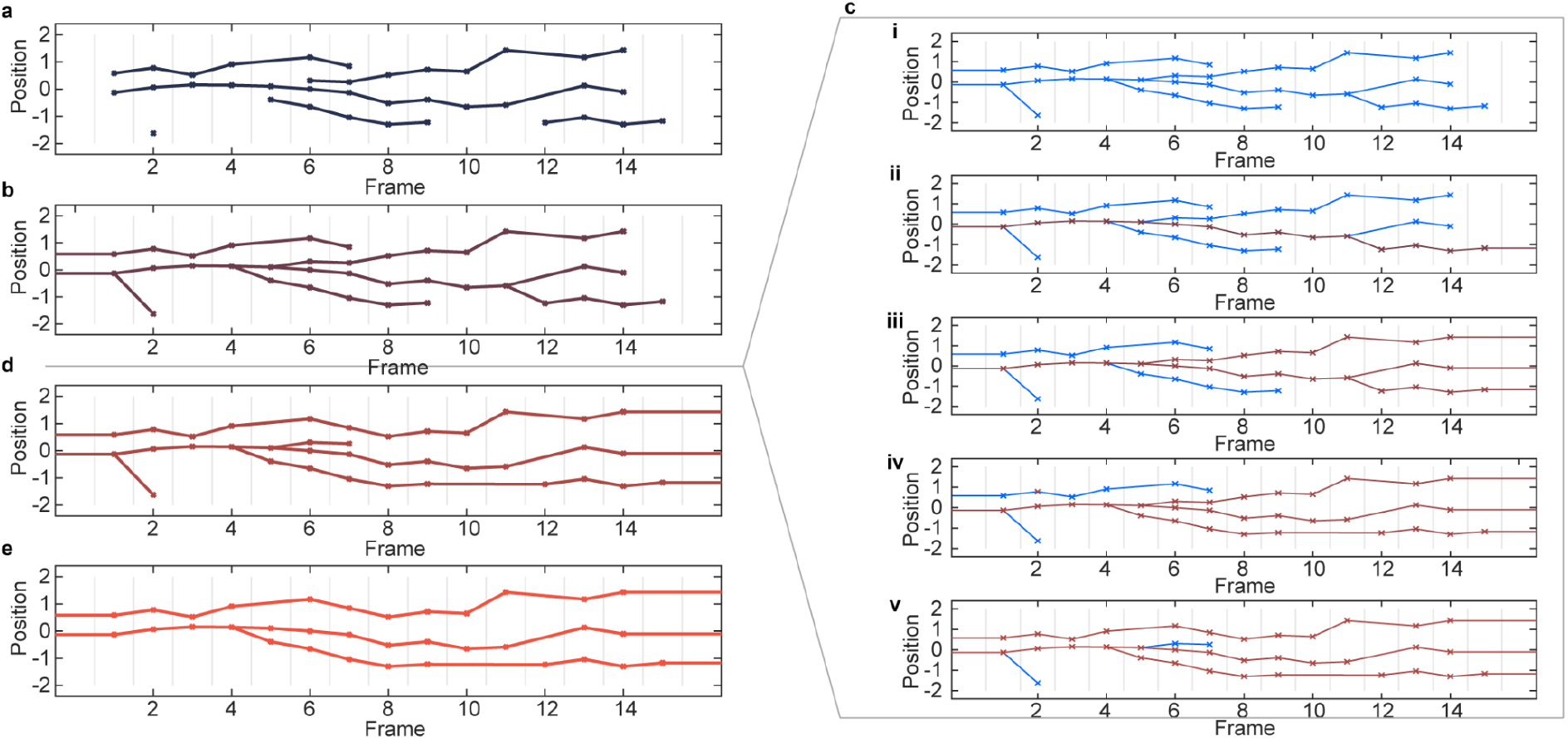
Stitching constrained A* output together. **a**, Example output of linked foci produced by constrained A* tracking. **b**, The first focus of every trajectory segment is linked to an earlier focus (chosen by lowest cost). Foci on the first frame are linked to imaginary foci before the first frame. **c**, Detailed overview of the procedure between **b** and **d**. **i**, The goal is to have as many foci as possible connected to an imaginary frame before the beginning (frame 0) and to an imaginary frame after the end (in this case frame 16) through its links to other foci. Initially no focus is connected in this way and we colour the entire tracking blue. **ii**, For each loose end of the tracking, starting at the last loose end (here on frame 15), we attempt to integrate it into the tracking by linking it to the imaginary last frame or replacing an existing link such that the overall tracking improves (see methods for details of how the cost of the entire tracking is calculated). Here the last loose end is linked to the imaginary last frame of the tracking. This results in 16 foci being connected as required (coloured red). **iii**, Same as in **ii** for the two loose ends on frame 14. 10 additional foci are now connected as required. **iv**, The next earliest loose end (frame 9) is interwoven by replacing the link from frame 11 to 12 by a link from frame 9 to 12. This results in 5 additional foci being connected. **v**, The next loose end (frame 7) is interwoven into the tracking by replacing an existing link. This connects 6 more foci, however since the replaced link was not part of a splitting event this also disconnects 2 foci from the imaginary last frame. The remaining two loose ends cannot be interwoven into the tracking in such a way that the tracking is improved. **d**, The result of the interweaving procedure. **e**, Remaining unconnected foci are labelled as false positives and removed.

**Fig. S4:**
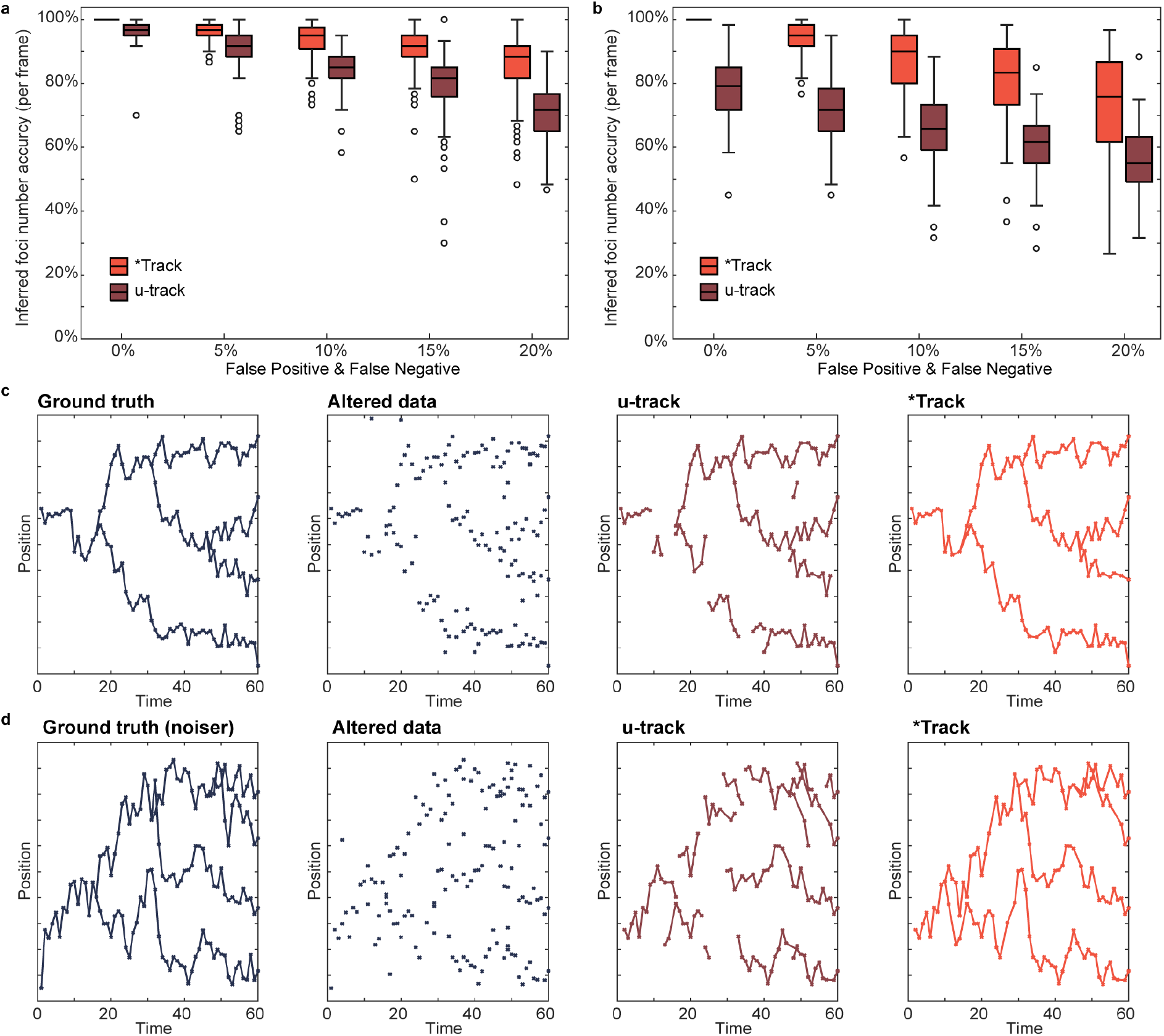
A comparison of ^★^Track and u-track. Due to the differences in the two methods (u-track can produce disconnected segments of the trajectory whereas ^★^Track cannot), we use the percentage of frames in which the number of inferred foci is equal to the ground truth as a measure of comparison of the methods. The parameters for u-track were chosen to give the best results and are found in Table S1 and in the code repository. **a**, Box charts comparing the accuracy of ^★^Track to u-track. The ground truth was altered similarly to Fig. 1c before being processed by the tracking algorithms. Each box chart was produced from 100 simulations. **b**, as in **a** but for more mobile foci as in Fig. 1d. **c** Example from **a** with 10% false positives & false negatives. **d**, Example from **b** with 10% false positives & false negatives.

**Fig. S5.**
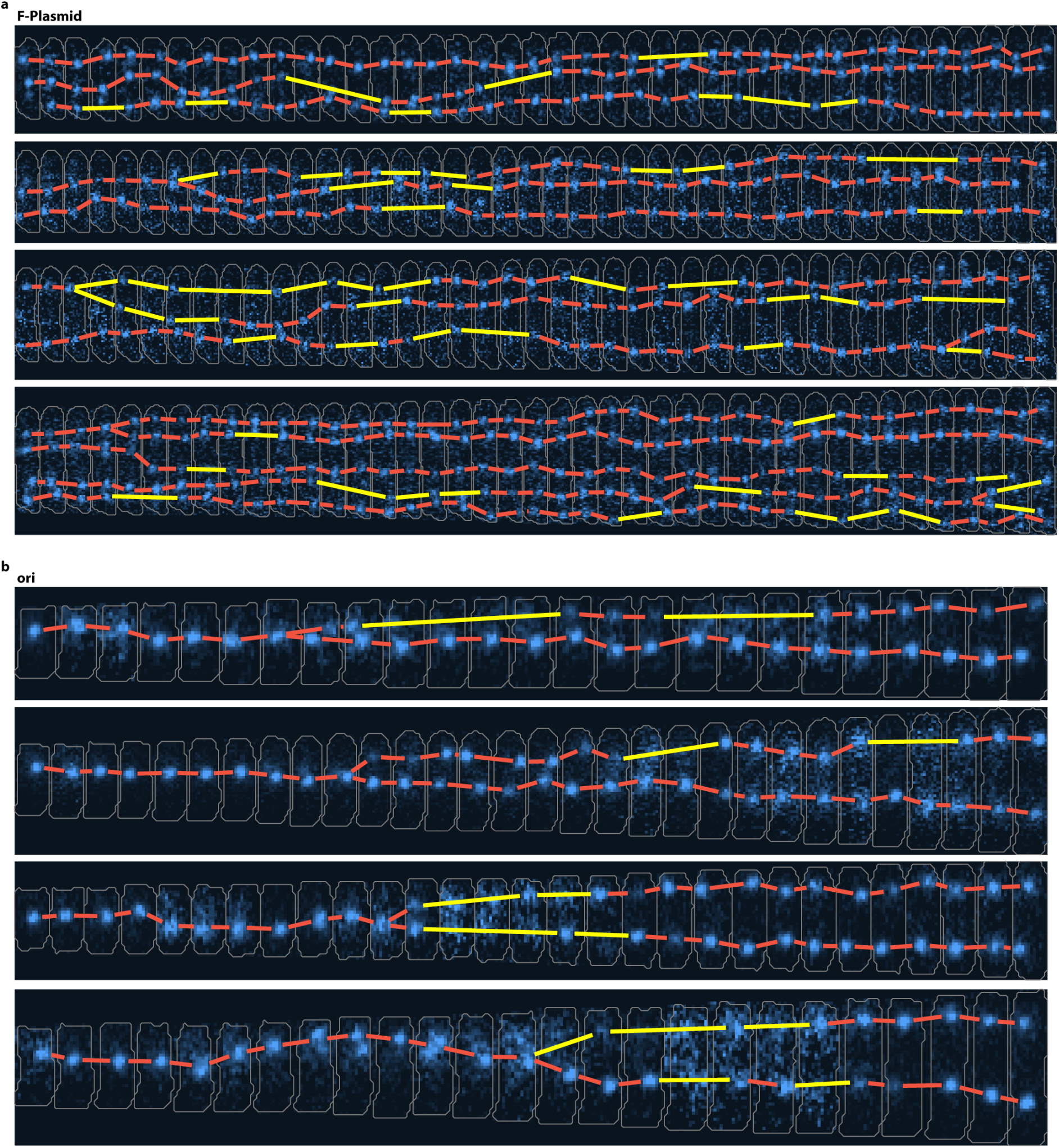
Examples of ^★^Tracked foci for ParB-mVenus labelled F-Plasmid and mTurqouise2-ParB labelled ori. **a**, cell cycle fragments (40 frames) from *E. coli* strain DLT3145 with tracked ParB-mVenus foci. Frame rate is 1 min^−1^. **b**, Cell cycles from *E. coli* strain IS130 with tracked mTurqouise2-ParB foci. Frame interval is 5 min. Yellow lines indicate links with frame skips.

**Fig. S6:**
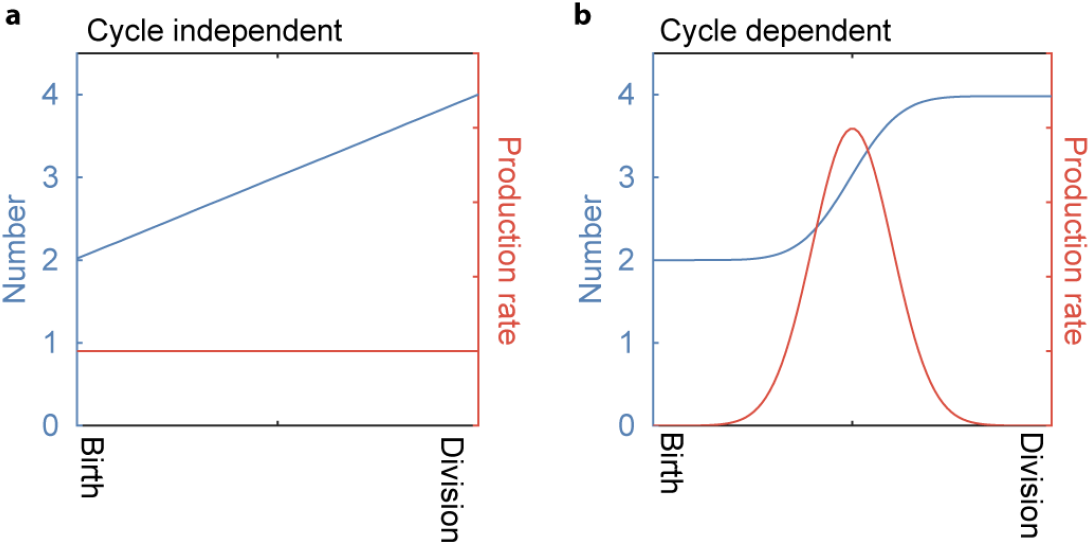
Cycle independent and cycle dependent replication. **a**, Cycle independent replication. At each point in time there is the same probability of replication (orange line). This causes the number (blue line) to increase linear **b**, Cycle dependent replication. There is a period during the middle of the cycle where replication occurs (orange line). This creates a sigmoidal change in the number.

**Fig. S7:**
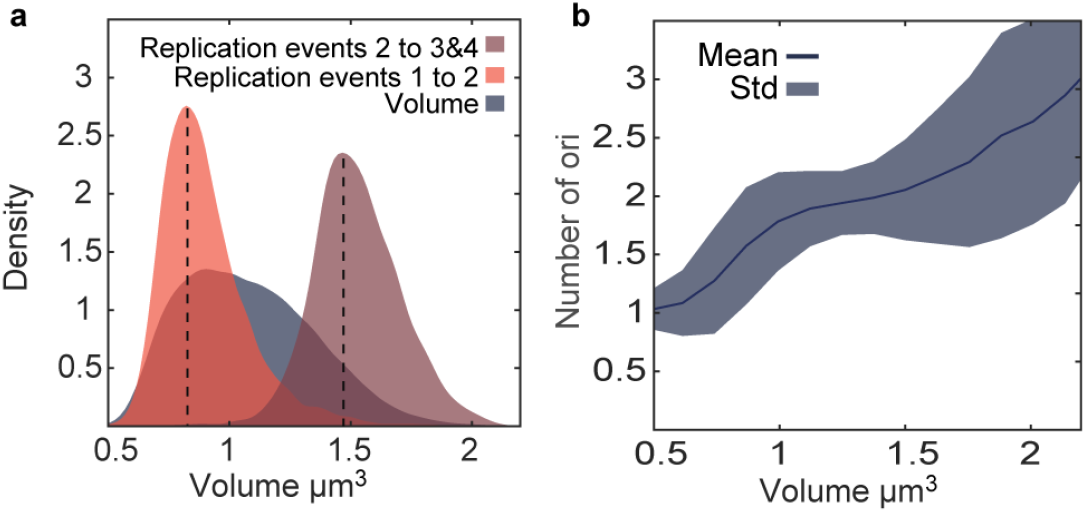
Second replication during the late cell cycle supports that volume per ori triggers replication. **a**, The distribution of replication events from 1 to 2 ori (orange) and from 2 to 3 or 4 ori (brown) plotted against cell volume. Black dashed lines indicate the peaks of the distribution at roughly 0.8 μm^3^ and 1.5 μm^3^. For reference the distribution of all measured volumes is plotted in gray. **b**, Mean number of ori ± standard deviation plotted against volume.

**Table S1.** Parameters used with u-track

| Parameter | Value |
| --- | --- |
| timeWindow | 6 |
| minTrackLen | 2 |
| linearMotion | 0 |
| minSearchRadius | 6 |
| maxSearchRadius | 6 |
| brownStdMult | 3 |
| useLocalDensity | 1 |
| brownScaling | [0.05, 0.01] |
| lenForClassify | 5 |
| linStdMult | 3 |
| linScaling | [1, 0.01] |
| maxAngleVV | 30 |
| gapPenalty | 5 |
| gapExcludeMS | 1 |
| strategyBD | 0.001 |

## Supplementary Videos

Supplementary Video 1: Video of the generation of the layered graph displayed in Fig. S2b. The links corresponding to the current path are shown in the bottom-middle.

Supplementary Video 2: Video of the A*-like path finding process. Only the colored part of the graph is generated. The links corresponding to the current best candidate path are shown in the bottom-middle. Additionally the partial cost of the edges of the candidate path, the heuristic estimate of completing this path and the total cost estimate are shown.

## Footnotes

1 As well as referring to the A* algorithm, we are inspired by the quote of Captain Kirk from Star Trek: *“There’s no such thing as the unknown, only things temporarily hidden”*.

